# Cortical interneurons ensure maintenance of frequency tuning following adaptation

**DOI:** 10.1101/172338

**Authors:** Ryan G. Natan, Winnie Rao, Maria N. Geffen

## Abstract

Neurons throughout the sensory pathway are tuned to specific aspects of stimuli. This selectivity is shaped by feedforward and recurrent excitatory-inhibitory interactions. In the auditory cortex (AC), two large classes of interneurons, parvalbumin- (PVs) and somatostatin- positive (SOMs) interneurons, differentially modulate frequency-dependent responses across the frequency response function of excitatory neurons. At the same time, the responsiveness of neurons in AC to sounds is dependent on the temporal context, with the majority of neurons exhibiting adaptation to repeated sounds. Here, we asked whether and how inhibitory neurons shape the frequency response function of excitatory neurons as a function of adaptation to temporal repetition of tones. The effects of suppressing both SOMs and PVs diverged for responses to preferred versus non-preferred frequencies following adaptation. Prior to adaptation, suppressing either SOM or PV inhibition drove both increases and decreases in spiking activity among cortical neurons. After adaptation, suppressing SOM activity caused predominantly disinhibitory effects, whereas suppressing PV activity still evoked bi-directional changes. SOM, but not PV-driven inhibition dynamically modulated frequency tuning as a function of adaptation. Additionally, testing across frequency tuning revealed that, unlike PVs, SOM-driven inhibition exhibited gain-like increases reflective of adaptation. Our findings suggest that distinct cortical interneurons differentially shape tuning to sensory stimuli across the neuronal receptive field, maintaining frequency selectivity of excitatory neurons during adaptation.

## Introduction

Neurons throughout the sensory pathway are tuned to specific aspects of stimuli, such as position and edge orientation in the visual cortex, or frequency and its modulation in the auditory cortex. This selectivity is thought to determine behavioral and perceptual discrimination in the natural world. Excitatory neuron selectivity is shaped by both feedback and recurrent networks, including excitatory-inhibitory interactions. These interactions are complex: inhibitory neurons exhibit remarkable diversity in their morphology and physiological properties, as well as the complexity in connectivity patterns, targeting excitatory as well as other inhibitory neurons (1–4). The two most common classes of inhibitory neuron, parvalbumin- (PVs) and somatostatin- (SOMs) positive interneurons, are thought to differentially shape excitatory neuronal responses. PVs predominantly target the cell bodies of excitatory neurons (5–8), whereas the majority of SOMs target the distal dendrites of excitatory neurons (9–12). In the auditory cortex, recent studies found that PVs and SOMs contribute to sound representation, tone frequency representation and behavioral selectivity to the stimuli (13–19).

Yet, whereas the majority of these studies relied on static stimuli, presented in isolation, auditory processing is inherently dynamic (20), and neurons throughout the auditory pathway, including the auditory cortex, adjust their response properties to sounds depending on the stimulus statistics. The most prevalent form of such changes occurs through adaptation. By reducing the signaling strength in response to commonly encountered stimuli, neuronal populations enhance sensitivity to changes in those stimuli (21). This process is thought to increase information transmission and computational efficiency of neuronal networks (22, 23). Adaptation is thought to rely largely on feedforward synaptic depression that drives gain adaptation (24). However, it can also be produced through recurrent network dynamics and, as shown in other modalities, recurrent inhibition can contribute to neuronal response dynamics over variable time scales (21, 25). Neurons in the auditory cortex exhibit selectivity to sound that is highly context-dependent: neurons throughout the auditory pathway, including those in the cortex, exhibit adapting responses depending on the temporal regularity of sounds. Recently, we and others showed that PVs and SOMs both provide enhanced inhibition in the adapted regime (19, 26). However, we currently lack an understanding of whether the differential action of PVs and SOMs generalizes to adaptation in all forms, and how PVs and SOMs transform tone-frequency representation in the auditory cortex before and after adaptation. Our goal was to arrive at an understanding of whether interneurons provide excitatory neurons with static or dynamic modulation across their frequency tuning curve and over time.

To investigate how PVs and SOMs differentially shape the tuning curve of excitatory neurons before and after adaptation, we presented repeated tones at different frequencies to an awake, head-fixed mouse, and recorded neuronal activity, while optogenetically inactivating either PVs or SOMs on a subset of trials (Figures 1 and 2).

**Figure 1.**
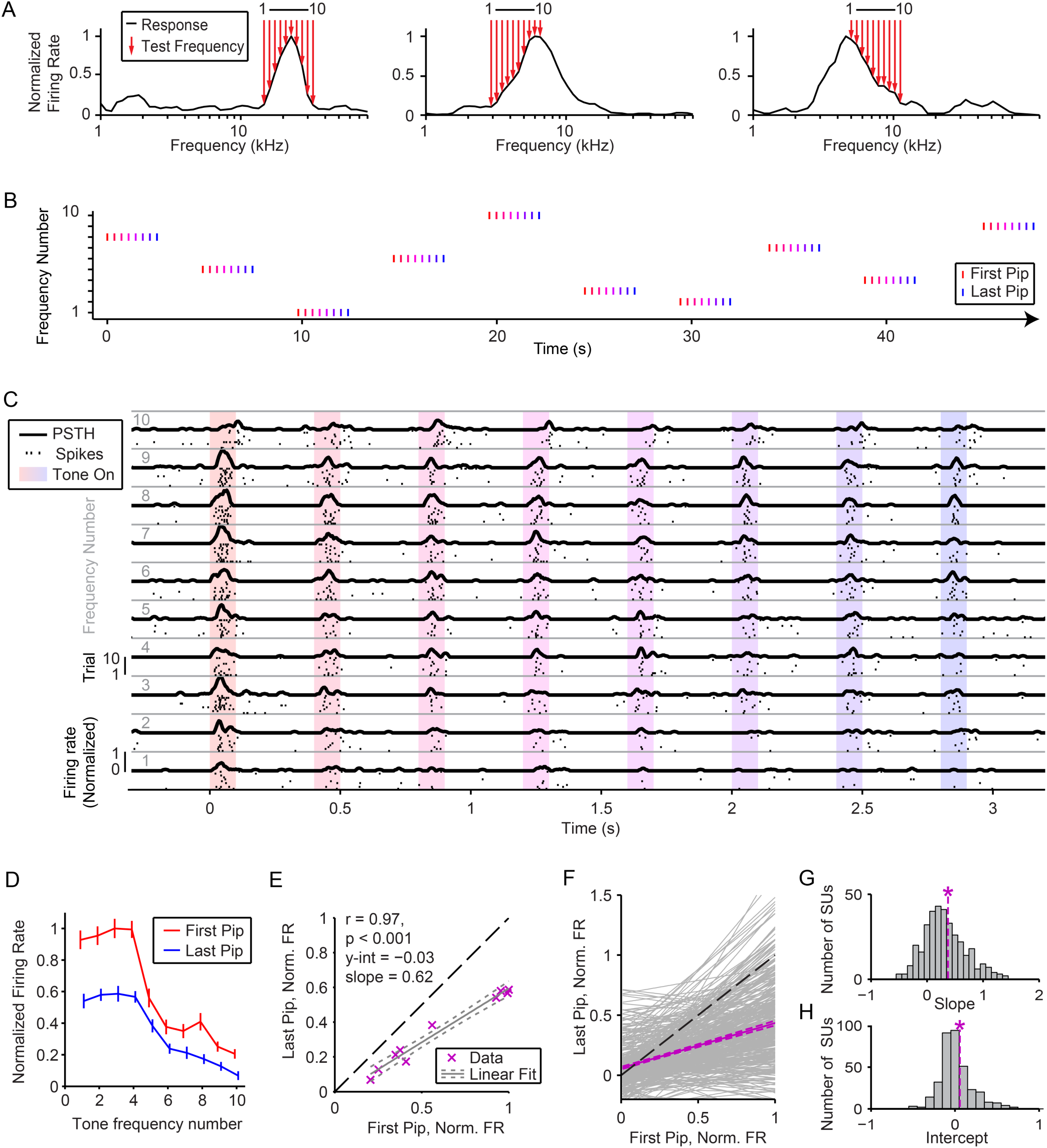
Adaptation scales responses across tuning curve. A) Frequency response functions (black line) for three neurons and the corresponding 10 evenly log frequency-spaced tones (red arrows) selected to construct the tone train stimulus set. B) Each tone train is composed of 8 tone pips of a single frequency. Each train is separated by a 1.6 seconds inter-trial interval, and tone frequency is selected in pseudorandom counterbalanced sequence. Tone mark color (red to blue) illustrates the progression of the train’s selected frequency from novel (red) to standard (blue). C) Raster plots and PSTHs depicting a single neuron’s response to tone trains of each frequency. Shaded areas indicate tone pips. Color as in B. D) Normalized mean firing rate of the same neuron in C in response to first (red) and last (blue) tone pip of each train across ten frequencies. E) Normalized mean firing rate of the same neuron in C in response to the first versus last tone pip across each of ten frequencies (purple ‘x’s). Grey lines - linear fit (solid) and fit error (dashed). Dashed black line – unity line. F) Linear fits to first versus last pip normalized firing rate responses for each neuron (grey) and population average (purple) pooled across SOM-Cre and PV-Cre. G-H) Slope (G) and intercept (H) of the linear fit for all neurons. Dashed line – population mean. Asterisk – population significantly different than 0 (intercept) or 1 (slope).

**Figure 2.**
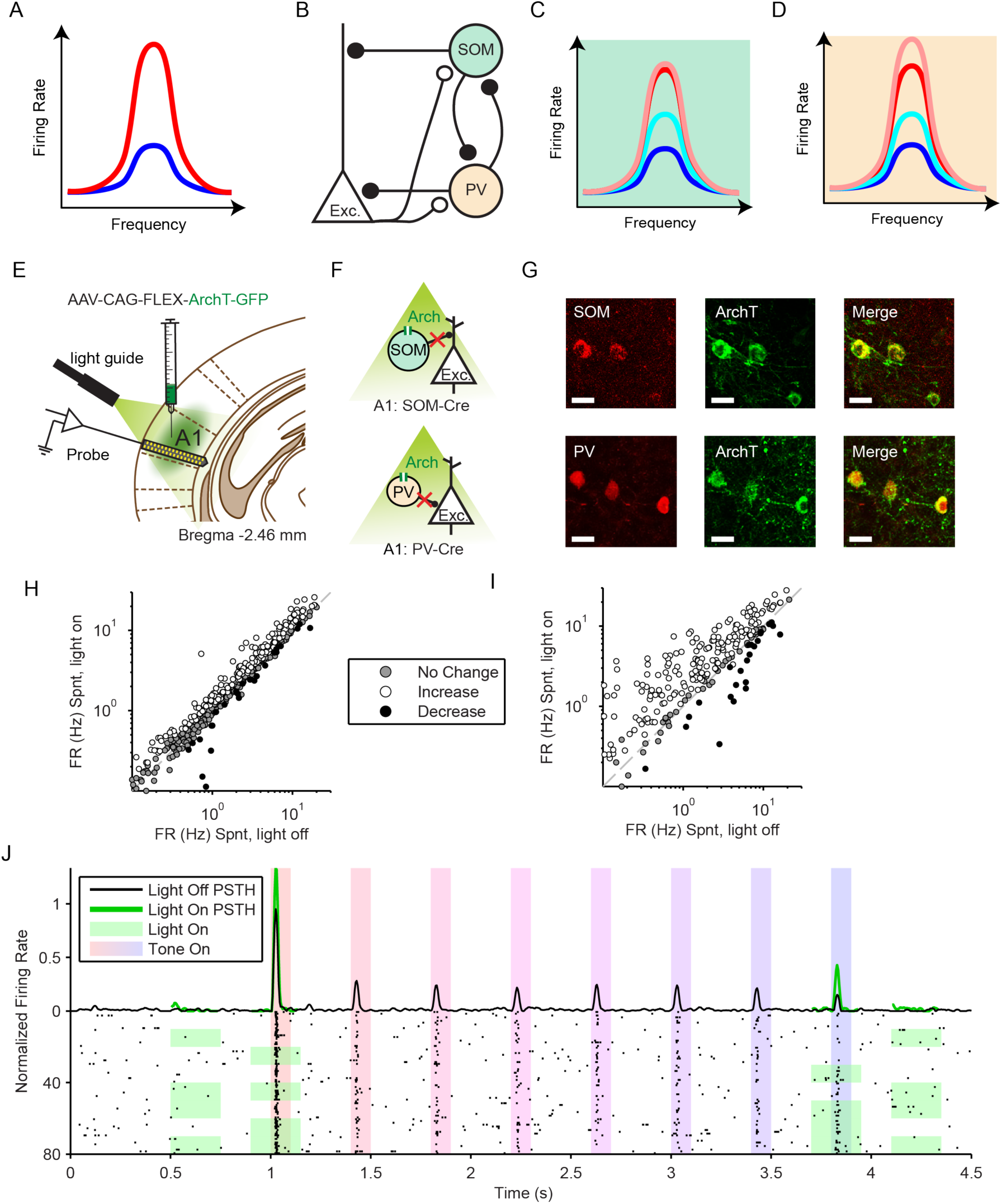
Experimental design for testing the function of interneurons in adaptation to repeated tones. A) Diagram of a cortical neuron frequency response function before adaptation (Red line) and predicted response after adaptation (Blue line). B) Diagram of key elements of cortical excitatory-inhibitory circuits studied. Somatostatin-positive interneurons (SOMs) make inhibitory synapses onto the distal dendrites of pyramidal neurons (Pyr) and onto parvalbumin-positive interneurons (PVs). PVs form inhibitory synapses onto proximal dendrites and somas of Pyrs and onto SOMs. Pyrs form excitatory synapses onto local SOMs and parvalbumin-positive PVs. C) Model of predicted modulatory effects of SOMs on Pyr tuning curve. Before adaptation, suppressing SOMs does not change responses across the tuning curve (pink line). After adaptation, suppressing SOMs increases responses across the tuning curve (light blue line). D) Model of predicted modulatory effects of PVs on Pyr tuning curve before and after adaptation. Before and after adaptation, suppressing PVs increases responses across the tuning curve. E) Diagram of optogenetic manipulations. AAV-CAG-FLEX-ArchT-GFP was injected in A1. During experiments, an optic fiber was positioned to target A1 and neuronal activity was recorded using a multi-channel silicon probe in A1. Bottom: Green light (532 nm) suppresses PVs in PV-Cre mice or SOMs in SOM-Cre mice. F) Green light (532 nm) suppresses SOMs in SOM-Cre mice (top) or PVs in PV-Cre mice (bottom). G) Transfection of interneurons with ArchT. Immunohistochemistry demonstrating co-expression of ArchT and an interneuron-type reporter in A1. Bottom: SOM-Cre mouse A1. Red: anti-body stain for somatostatin. Green: Arch-GFP. Merge; co-expression of ArchT and somatostatin. Top: PV-Cre mouse A1. Red: antibody stain for parvalbumin. Green: Arch-GFP. Merge; co-expression of ArchT and parvalbumin. Scale Bar = 25 µm. H and I) Spontaneous firing rate with versus without optogenetic suppression of SOMs (H) or PVs (I). Each dot represents a single neuron and indicates that optogenetic suppression significantly increased (white), decreased (black) or had no effect on spontaneous firing. J) Raster plot and PSTH depicting of a single neuron’s spiking responses with (green) and without optogenetic modulation (black). Green shading in raster plot indicates light pulse times and trials.

## Results

### Cortical neurons exhibit adaptation to the stimulus

To understand whether and how frequency tuning of a neuron influences its adaptation to a repeated tone, we first measured neuronal frequency response functions across a broad range and then presented repeated tones across a limited frequency range (Figure 1A, B) in awake, head-fixed mice. As illustrated by three representative neurons, the ten chosen tone frequencies covered a range of the neuronal frequency response function that evoked different response amplitudes (Figure 1A). Since many neurons were tested simultaneously, the chosen test frequencies effectively sampled different portions of the tuning curve for different neurons in each experimental session. Each 100ms tone was repeated 8 times at 2.5 Hz, followed by 2.4 s of silence to allow adaptation to reverse (Figure 1B). Whereas cortical neurons adapt across a range of tone repeat rates (27), a relatively long (300 ms) inter-tone interval was chosen to incorporate the timecourse of optogenetic stimulation and to enable comparison of results to prior studies (19). Furthermore, this timecourse was expected to target long-term adaptation, likely affected by intracortical feedback mechanisms which take place over hundreds of milliseconds (19, 28). Neurons responded to repeated tones with an initially strong response, which gradually reduced over tone pip repeats (Figure 1C), exhibiting adaptation.

### Temporal adaptation scales divisively neuronal responses across the tuning curve

We first assayed the structure of adaptation to repeated tones across neuronal tuning curves. The degree to which neuronal tone-evoked responses adapt can be dependent upon frequency selectivity. If adaptation corresponds to a subtractive effect affect, we expect the change in the neuronal firing rate to be reduced by a constant amplitude across the frequency response function, whereas if adaptation is divisive, the reduction in firing rate would scale with the neuron’s responsiveness to tones. Understanding whether an effect is subtractive or divisive can give hints about the biophysical basis for its mechanism – e.g. suppression of excitatory neuronal responsiveness due to distal dendrite inhibition is expected to produce a linear, rather than a multiplicative shift in responses; whereas suppression closer to the cell body might be expected to produce a multiplicative scaling effect (16).

We measured the mean spiking response to the first and last tone pip at each of 10 frequencies (Figure 1D). For each neuron, plotting the first tone response to each test frequency on the x-axis sorts the neuron’s tuned responses from preferred to non-preferred frequency, and allows us to model tuning before and after adaptation (Figure 1E). In an adapting neuron, a linear fit of the tone-evoked neuronal responses to the paired first versus last tone across frequencies reveals a non-significant y-intercept, and a significant slope <1. These parameters indicate divisive scaling. For all neurons that exhibited any optogenetic modulation during tone-evoked activity, in either SOM-Arch (n = 184 neurons from 6 mice over 4 sessions each) or PV-Arch (n = 169 neurons from 5 mice over 4 sessions each) groups, the average responses exhibited a divisive adaptation effect with little evidence for subtraction (slope < 1, and intercept not significantly different from 0) (Figure 1F-H). Over individual neurons, however, there was a mix of divisive and subtractive effects. Some units exhibited a y-intercept significantly different than 0, pointing to a linear shift. Most units had either a positive or a negative slope with amplitude less than 1, suggesting the adaptation/facilitation was typically stronger for tones at preferred than non-preferred frequencies. These findings overall support a *divisive adaptation* model for temporal adaptation to repeated sounds.

### Optogenetic manipulation of PV and SOM activity

Divisive adaptation requires that the response attenuation across the tuning curve be frequency specific, i.e. spiking is reduced more strongly in the center as compared to the tuning sidebands (Figure 2A). Following on our recent finding for differential control of SSA by PVs and SOMs, we hypothesized that these neurons may play distinct roles in temporal adaptation: PVs would provide uniform suppression in both adapted and non-adapted states (Figure 2C), whereas SOM suppression would be selective for the adapted state, and be greater for frequencies in the center of the receptive field and weaker for receptive field sidebands (Figure 2D).

To test this hypothesis, we used viral transfection to drive selective expression of Archaerhodopsin in SOMs or PVs in the auditory cortex of SOM-Cre or PV-Cre mice, respectively (Figure 2E, F), in order to suppress the activity of the respective interneuron type upon presentation of light over the auditory cortex. Viral expression was confirmed post-mortem via immunohistochemistry (Figure 2G) consistent with previous results in our laboratory (18, 19). Illuminating the auditory cortex increased neuronal activity in many recorded neurons, as expected due to the optogenetic suppression of inhibitory activity in either SOM-Arch or PV-Arch mice (Figure H, I). In SOM-Cre mice, 42% of neurons exhibited increased, and 5% of neurons exhibited decreased spontaneous activity, whereas 53% were not significantly affected by the optogenetic manipulation (Figure 2H). In PV-Cre mice, 74% of neurons exhibited increased, and 9% of neurons exhibited decreased spontaneous activity, whereas 17% were not significantly affected by the optogenetic manipulation (Figure 2I). The neurons that exhibited significantly suppressed firing rate due to manipulation were considered putative Cre-expressing interneurons, and were excluded from further analysis of the putative pyramidal neuron population.

The laser was presented on half of all trials either during the first or last tones within each block, and either preceding the first tone, or following the last tone in a block-randomized fashion (Figure 2J). Laser presentation affected differential baseline and tone-evoked activity. For a representative neuron, the spontaneous firing rate was only slightly increased (Figure 2J) either before or after the tone train during light-on trials, whereas the responses to the first and the last tone were significantly increased.

### The effects of SOM, but not PV suppression, differ between adapted and non-adapted regimes

We hypothesized that the effects of suppressing SOM and PV activity differed across the neuronal tuning curve with adaptation. In order to test how SOM and PV activity impacted tuned responses relative to adaptation, we selected a subset of putative excitatory neurons that individually exhibited frequency tuning both before and after adaptation, i.e. each neuron’s first vs. last tone response profile (as in Figure 1E) exhibited slope greater than 1, and intercept was not significantly different from 0 (Supplementary Figure 1A, B). These subsets of neurons were well suited to represent the range of frequency dependent responses because stimulus test frequencies spanned each neuron’s frequency response function from near best frequency (within 2/5ths of an octave) to below the half-maximum response strength. We first confirmed that adaptation was similar in neurons from SOM-Cre (n = 55) and PV-Cre (n = 23) groups: there was no significant difference in the slope (p = 0.211, t(76) = 1.26) or intercept (p = 0.714, t(76) = 0.37) (Supplementary Figure 1C, D), and the population mean responses reflected divisive adaptation similar to the whole population (as in Figure 1F). As expected, each group’s population mean neuronal response to tone trains exhibited adaptation (Figure 3A, B), responding to first tone pips more strongly than to last tone pips. Typically, neuronal firing rates were disinhibited when either PVs or SOMs were suppressed with light before or after the tone train.

**Figure 3.**
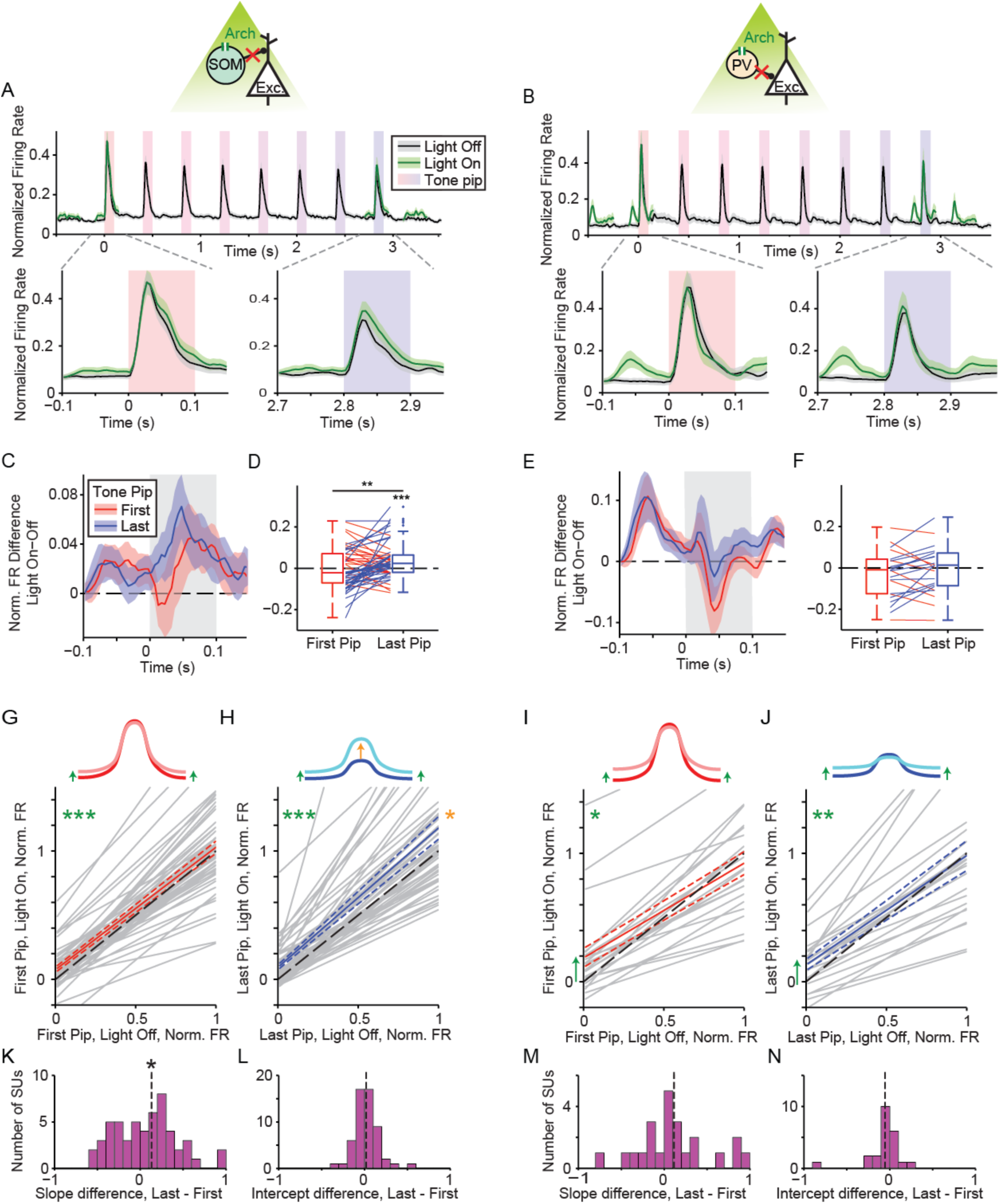
SOM inhibition increases with stimulus repetition and contributes to scaling after adaptation. A and B) Selected neurons (Fig. 3 F and G) population average firing rate in response to tone trains with (green) and without (black) optogenetic suppression of SOMs (A) and PVs (B). Top: PSTH over whole tone train. Bottom: PSTH during first (left) and last (right) pip. C and E) Overlay of PSTHs of the mean per-neuron difference in firing rate between trials with and without optogenetic suppression of SOMs (C) and PVs (E) for the first (red) and last (blue) pip. D and F) Summary of per-neuron differences in firing rate between trials with and without optogenetic suppression in the first and last tone onset response (0-50ms from tone onset) for SOMs (D) and PVs (F). Line color indicates that the effect of suppression increased (blue) or decreased (red) from the first to last pip. G-J) Top: Model depiction of tuning curve in response to the first (red) and last (blue) tone pip with (light) and without (dark) optogenetic suppression of SOMs (G and H) or PVs (I and J). Arrows emphasize significant modulation for non-preferred tones (green) and preferred tones (Orange). Bottom: Linear fits to the first (G and I) or last (H and J) tone pip firing rate responses in selected neurons, with versus without optogenetic suppression of SOMs (G and H) or PVs (I and J) for each neuron (grey) and population average (red or blue). Green asterisks – significant changes in non-preferred frequencies. Orange asterisks – significant changes in preferred frequencies. K-N) Slope (K and M) and intercept (L and N) of the linear fit for selected neurons in SOM-Cre mice (K and L) or PV-Cre mice (M and N), respectively. Red - first pip. Blue - last pip. Dashed line – mean value.

Suppressing SOM neurons drove differential disinhibition, specific to the adapted state. SOM suppression had no overall effect on responses to the first tone pips over the recorded population (p = 0.985, t(54) = -0.02), but significantly disinhibited responses to the last tone pips (p = 0.001, t(54) = 3.40). Change in neuronal responses due to SOM inhibition significantly increased from the first to last tone pip (p = 0.007, t(54) = 2.79) (Figure 3C, D). The time course of the difference in the tone-evoked responses during light-on and light-off conditions differed significantly between responses to the first and last tone pips, further illustrating that adaptation shifted the effect of SOM suppression toward disinhibition of excitatory neurons. These findings suggest the SOMs provides tone-evoked inhibition, which increases in the adapted regime.

In contrast, whereas suppressing PVs drove a significant increase in spontaneous activity, PV suppression resulted in no significant change in the tone-evoked response over the neuronal population for either the first (p = 0.701, t(22) = -0.39) or last tones (p = 0.688, t(22) = 0.41) (Figure 3E, F). Suppressing PVs drove a similar amount of suppression and activation, resulting in a non-significant difference across the population and no significant difference between the adapted and non-adapted responses (p = 0.138, t(22) = 1.54). These results suggest that PVs provide equal amount of excitation and inhibition to the excitatory neurons in either adapted or non-adapted state.

### Effects of SOM, but not PV, inhibition become stronger with adaptation in a frequency-selective fashion

We next tested whether the effects of SOMs or PVs differed for tones in the center (preferred frequencies) and on the sidebands (non-preferred) of the tuning curve of excitatory neurons (Figure 3G, H, I, J). We compared the response change due to SOM or PV suppression on excitatory responses to tones across the entire tuning curve, for either the first (Figure 3G, I) or last tone (Figure 3H, J). For the first tone, suppressing SOMs slightly disinhibited responses only for non-preferred frequencies (non-preferred p = 2e-4, t(54) = 4.01; preferred p = 0.548, t(54) = 0.62). Yet in the adapted regime, suppressing SOMs preferentially disinhibited responses to tones at preferred frequencies (non-preferred p = 3e-6, t(54) = 5.26; preferred p = 0.045, t(54) = 2.05) (Figure 3G, H). Indeed, there was a significant positive shift in the firing rate for preferred frequencies in the suppression-versus-tone frequency curve (slope, p = 0.031, t(54) = 1.91) and no change in the intercept (p = 0.155, t(54) = 1.02). These results suggest that SOMs increasingly contribute to adaptation of responses across frequency tuning in a selective fashion (Figure 3K, L).

In contrast, suppressing PVs had a stronger disinhibitory effect on neuronal responses at non-preferred tone frequencies than at preferred frequencies during both the first (non-preferred p = 0.014, t(22) = 2.68; preferred p = 0.392, t(22) = -0.87) and last tone pips (non-preferred p = 0.004, t(22) = 3.26; preferred p = 0.880, t(22) = 0.015) (Figure 3I, J). Over the population, neither the slope (p = 0.103, t(22) = 1.31) or intercept (p = 0.894, t(22) = 1.29) significantly changed from the first to last tone (Figure 3M, N). This suggests that PVs preferentially inhibit neuronal responses in the side-bands of frequency curves, and their effect is insensitive to adaptation to tone repetition.

### Effects of SOM and PV suppression differ for adaptive and non-adaptive neurons

To examine how the effects of optogenetic manipulation of SOM or PV activity changed with adaptation for individual neurons, we compared the change in tone-evoked responses of excitatory neurons for the first and last tone among adapting and non-adapting neurons separately (Figure 4, Supplementary Figure 2). For the adapting population, neuron-frequency pairs for which the first tone pip evoked significantly stronger spiking than the last pip were included (Figure 4A, C, Supplementary Figure 2A, C). For the non-adapting response population, only neuron-frequency pairs were included from neurons in which the first and last tone pip-evoked spiking was not significantly different at any frequency (Figure 4B, D, Supplementary Figure 2B, D). We only detected a small number of neurons that exhibited FR facilitation, rather than adaptation, in response to tone repetition and excluded them from further analysis (2 neurons among PV-Cre and 5 among SOM-Cre populations).

**Figure 4.**
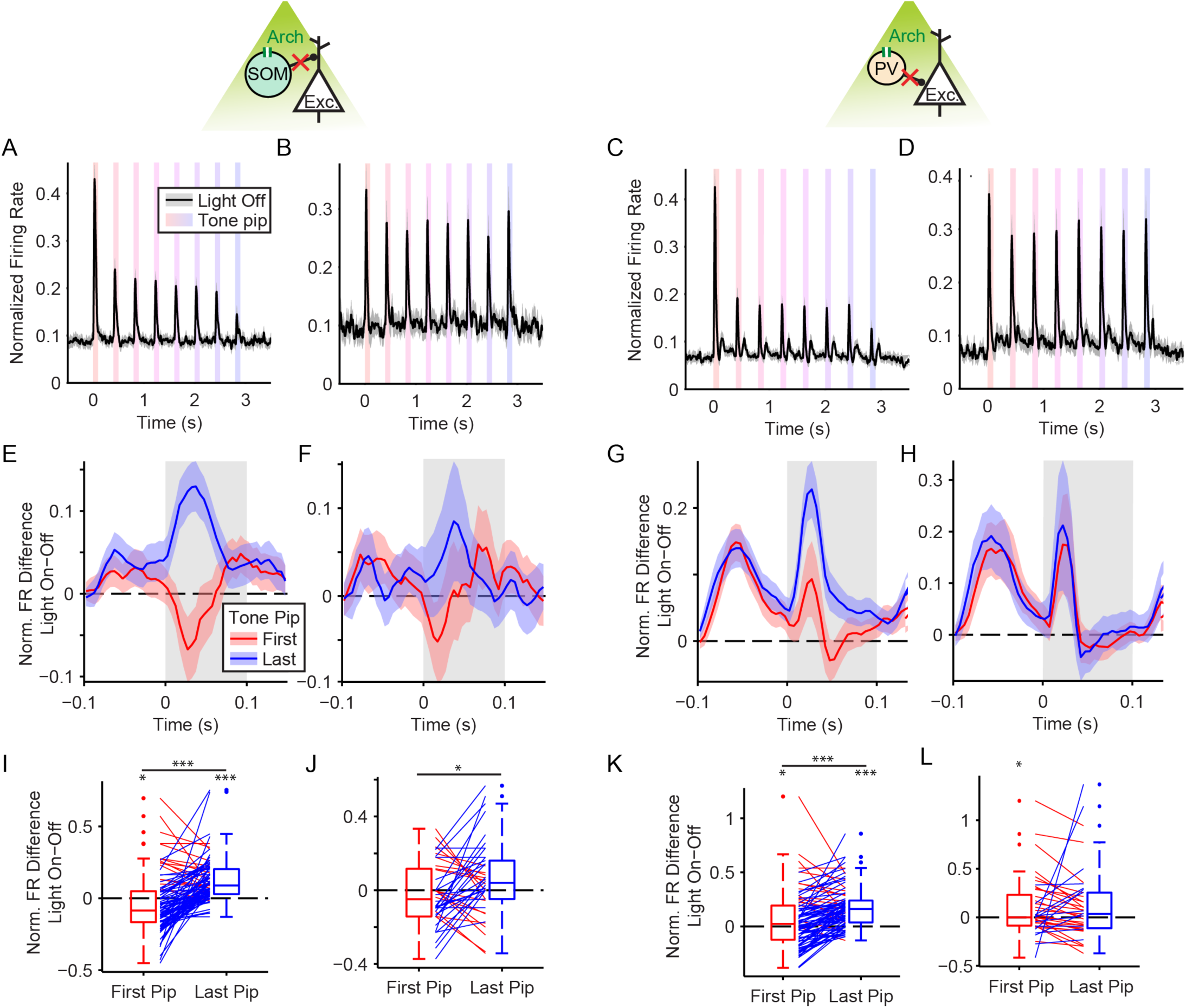
Effects of optogenetic modulation of SOMs and PVs on adaptive and non-adaptive neurons. A-D) Population average PSTH of normalized firing responses to tone trains for adaptive neuronal responses (A and C) and non-adaptive neuronal responses (B and D) among the SOM-Cre (A and B) or PV-Cre populations. E-H) Overlay PSTHs of the mean per-neuron difference in firing rate between trials with and without optogenetic suppression of interneurons for the first (red) and last (blue) pip. Panels E, F, G and H correspond with panels A, B, C and D, respectively. Grey shaded region indicates tone on time. For light-on trials, illumination lasted the entire duration of the PSTH window. I-L) Summary of the per-neuron difference in firing rate between trials with and without optogenetic suppression of interneurons in the first and last tone onset response (0-50ms from tone onset). Line color indicates that the effect of suppression increased (blue) or decreased (red) from the first to last pip. Panels I, J, K and L correspond with panels A, B, C and D, respectively.

SOM suppression affected first and last tone pip-evoked responses differentially between adaptive and non-adaptive neurons. Among adapting neurons, SOM suppression led to heterogeneous first tone pip-evoked modulation; 30% increased and 34% decreased (Figure 4E, Supplementary Figure 2E). By contrast, more last tone pip-evoked responses increased (54%) and only a small fraction decreased (2%) (Supplementary Figure 2I). Interestingly, on average, SOM suppression led to significant inhibition of first tone pip-evoked responses (p = 0.027, t(96) = -2.25) and significant disinhibition of last tone pip-evoked responses (p = 2e-12, t(96) = 8.04) (Figure 4I, Supplementary Figure 2 M, Q), showing that the strength of SOM inhibition increased from the first to the last tone pip (p = 2e-11, t(96) = 7.63), as already observed for the subpopulation of adapting tuned neurons (Figure 3G, H). Similar to adapting units, non-adapting units were heterogeneously modulated by SOM suppression during the first tone pip (31% increased, 28% decreased) (Figure 4J, Supplementary Figure 2B, F, N). Unlike adapting units, non-adapting units continued to respond heterogeneously during the last tone pip (38% increased, 17% decreased) (Figure 4J, Supplementary Figure 2J, R). On average, SOM suppression of non-adapting units evoked no significant change of first (p = 0.373, t(41) = -0.90) or last tone pip-evoked responses (p = 0.065, t(41) = 1.90), but the strength of SOM inhibition increased from the first to the last tone pip (p = 0.027, t(41) = 2.29) (Figure 4J).

Similar to SOMs, PV inhibition affected first and last tone pip-evoked responses differentially between adaptive and non-adaptive responses. However, the net effects of PV suppression were weaker than those of SOM suppression. Among adapting units, PV suppression led to heterogeneous first tone pip-evoked modulation (27% increased and 28% decreased) (Figure 4C, G, K, Supplementary Figure 2G). For the last tone pip, many responses increased (66%) and a smaller portion decreased (8%) (Supplementary Figure 2K). On average, PV suppression lead to significant disinhibition of both first (p = 0.045, t(88) = 2.03) and last tone pip-evoked responses (p = 1e-13, t(88) = 8.75) (Supplementary Figure 2O, S), and the strength of PV inhibition increased from the first to the last tone pip (p = 1e-7, t(88) = 5.72) (Figure 4G, K). Non-adapting units were heterogeneously modulated by PV suppression during the first tone pip (31% increased, 29% decreased) (Figure 4D, H, L, Supplementary Figure 2H), and inhibited more responses during the last tone pip (24% increased, 43% decreased) (Supplementary Figure 2L). On average, PV suppression led to just significant disinhibition of first tone pip-evoked responses (p = 0.043, t(40) = 2.09) and no significant modulation of last tone pip-evoked responses (p = 0.051, t(40) = 2.01) (Figure 4L, Supplementary Figure 2P, T). The strength of PV inhibition did not change significantly from the first to last tone pip (p = 0.778, t(40) = 0.28) (Figure 4L) for non-adapting units. Thus, PV suppression provided a differential, but weaker contribution to adaptation than SOM suppression.

### Adaptation strength of excitatory neurons correlates differentially with the PV and SOM modulatory effects

Does the effect of SOM or PV suppression predict the strength of adaptation? If inhibitory neurons contribute to adaptation, we expect the magnitude of adaptation to correlate with the strength of response modulation due to interneuron suppression. In order to test this, we measured the correlation between the magnitude of adaptation, indexed by firing rate before and after inhibition, with the inhibition strength, indexed by firing rate with and without interneuron suppression, for several frequencies across neurons. To compensate for correlations between multiple samples from each neuron, we used cluster-robust standard error to measure regression (29). Indeed, the magnitude of adaptation showed a nearly significant correlation with the modulation of the firing rate due to SOM suppression for the first tone (r = 0.31±0.18, p = 0.044, t(173) = 1.71, n = 330), and this correlation grew even stronger and more significant for the last tone (r = 0.58±0.08, p = 2e-11, t(173) = 7.03, n = 330) (Figure 5A, C). The strength of SOM inhibition of each neuron is thus predictive of magnitude of adaptation. By contrast, the magnitude of adaptation was anti-correlated with the modulation of the firing rate due to PV suppression for the first tone (r = -0.46±0.19, p = 0.008, t(149) = -2.46, n = 466) and not correlated with for the last tone (r = 0.14±0.15, p = 0.181, t(149) = 0.15, n = 466) (Figure 5B, D). The strength of FR changes from PV modulation is thus predictive of the magnitude of adaptation, yet suppressing PVs typically drives the firing rate in the opposite direction from SOM. Furthermore, adaptation abolishes this correlation. Together, these results show that inhibition from SOMs, rather than PVs, matches the range of adaptation profiles across the population.

**Figure 5.**
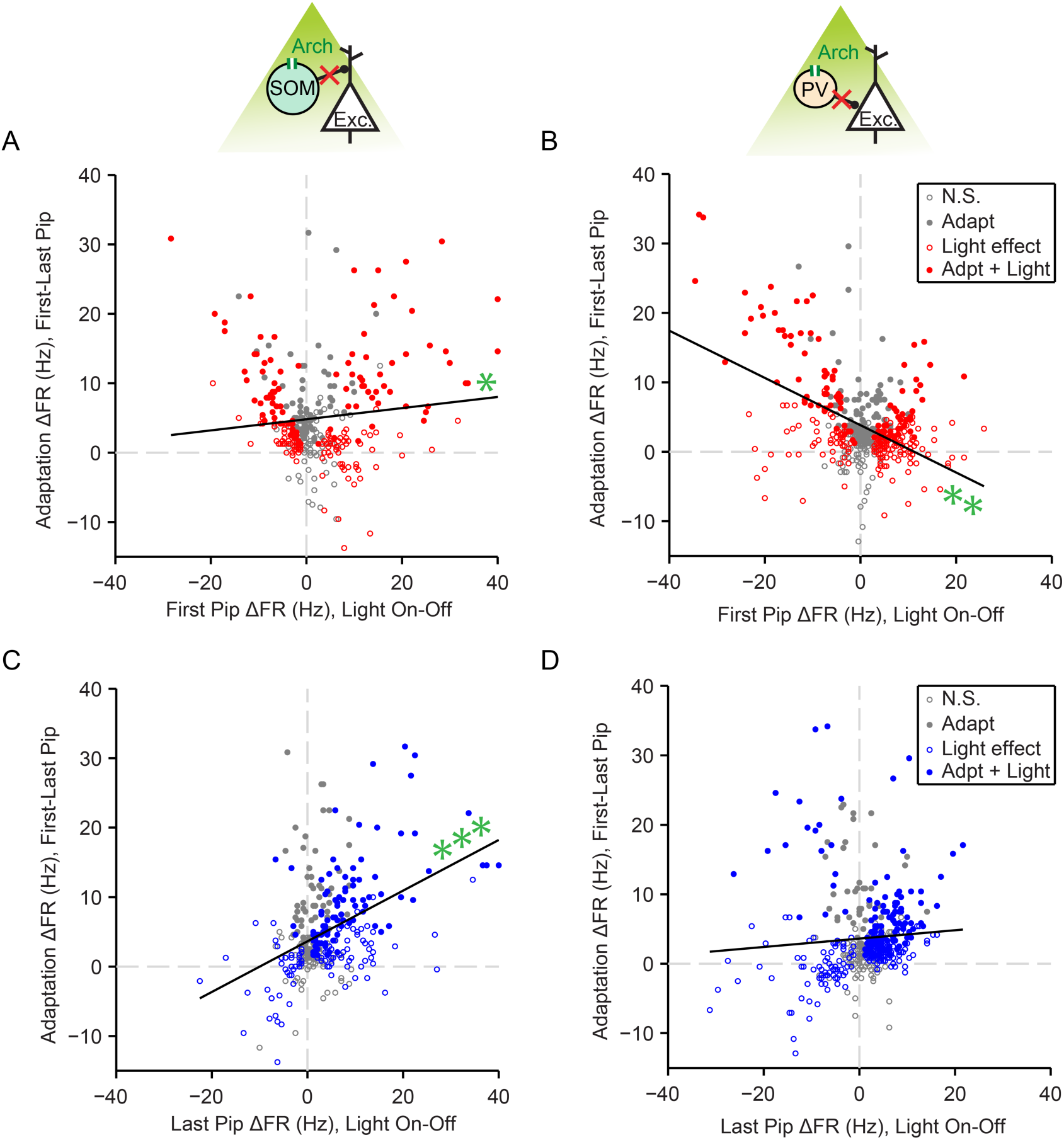
Strengths of SOM and PV inhibition differentially correlate with magnitude of adaptation. A-D) Effect of SOM (A and C) or PV (B and D) optogenetic suppression on firing rates for all neurons in response to the first (A and B) or last (C and D) tone pip versus the magnitude of firing rate adaptation. Optogenetic effects are measured as difference between the means of trials with and without optogenetic suppression. Magnitude of adaptation is measured as the difference between firing rate in response to the first versus last tone. Each dot represents a neuron-frequency pair. Dot color indicates that optogenetic suppression significantly modulated (red or blue) or did not modulate (grey) the responses to the tone pips, and filled dots indicate significant adaption. Black line: ordinary least squares fit. Green asterisks indicate the significance of correlation coefficient of cluster-robust regression.

Supporting these observations, the difference in the change in neuronal firing rate due to SOM suppression between during the first and last tone was not correlated for adapting neurons (r = 0.02±0.08, p = 0.0401, t(96) = 0.25, n = 161) or non-adapting neurons (r = 0.19±0.13, p = 0.075, t(40) = 1.47, n = 66) (Supplementary Figure 3A, C). These correlations demonstrate that the sign and magnitude of SOM inhibition can change over repeated stimulation, especially for neurons that significantly adapt. By contrast, the effect of PV suppression between the first and last tone was strongly correlated for both adapting units (r = 0.26±0.06, p = 1e-5, t(88) = 4.42, n = 186) and non-adapting units (r = 0.37±0.08, p = 4e-5, t(39) = 4.45, n = 101) (Supplementary Figure 3B, D). This result shows that the sign and magnitude of PV inhibition is largely insensitive to stimulus repetition or magnitude of adaptation. Together, these results show that neurons that experience stronger SOM inputs are more likely to adapt strongly, whereas modulation of activity by PVs is not affected by adaptation.

### Both SOMs and PVs affect excitatory neuronal activity more strongly after adaptation

We hypothesized that the observed effects could be explained by changes in connectivity strength between the inhibitory and excitatory neurons before and after adaptation. To approximate how strongly excitation of inhibitory neurons affects responses in excitatory neurons, we drove either SOMs or PVs to express an ultrafast Channelrhodopsin (ChETA) and used brief light pulses to elicit spiking in these interneurons with high temporal precision during tone pip trains or spontaneous activity (Figure 6A-D). Immunohistological verification of brain tissue identified co-localization of ChETA and either parvalbumin or somatostatin, confirming the specificity of expression of ChETA in interneurons (Supplementary Figure 4A). Brief pulses of light elicited reliable spikes within 3-7 ms of laser onset in a subset of neurons, indicating that light activated those neurons directly (Figure 6B, C; Supplementary Figure 4B-E).

**Figure 6.**
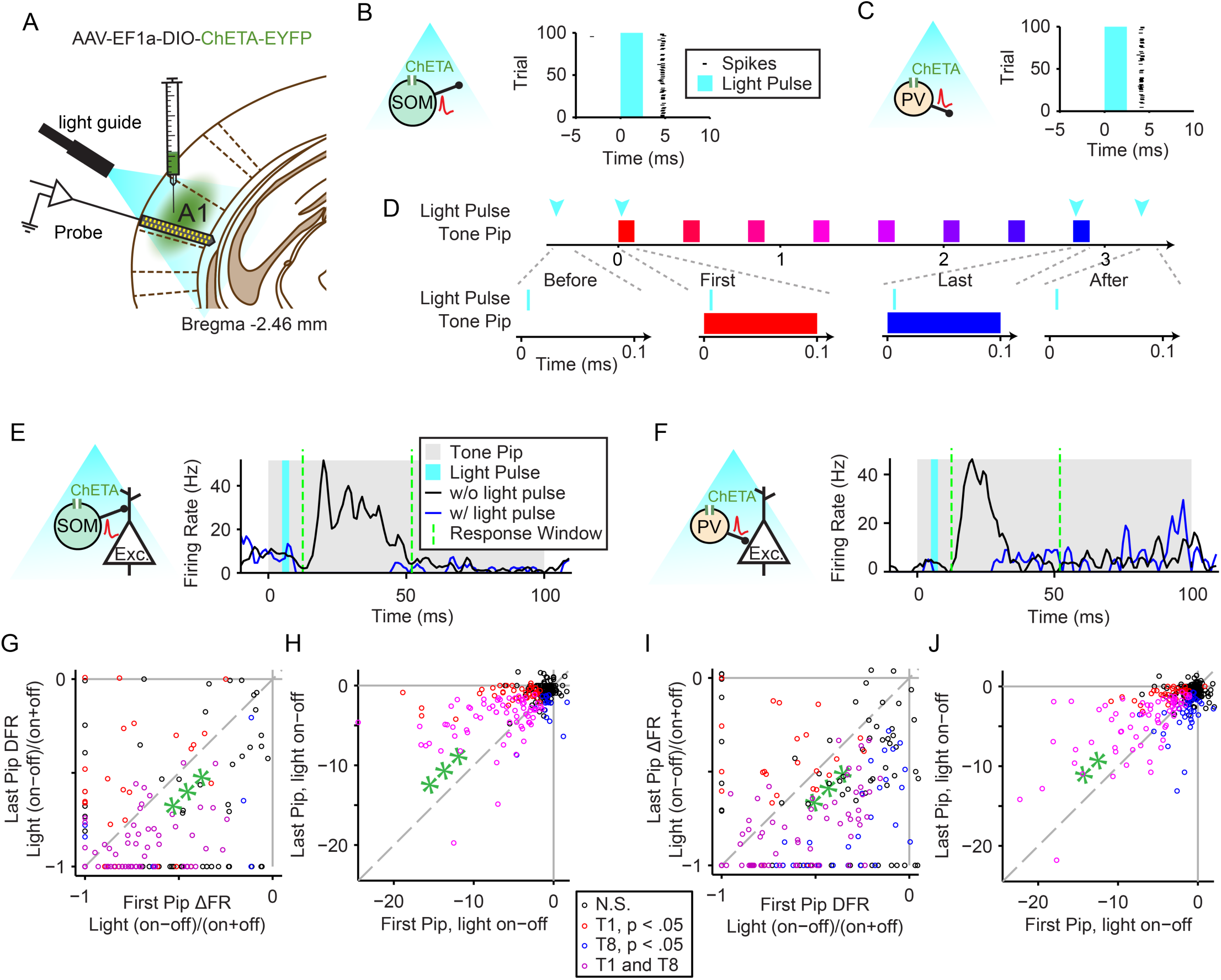
Effects of brief SOM or PV activation on tone-evoked responses before and after adaptation. A) Diagram of optogenetic methods. A1 was injected with AAV-EF1a-DIO-ChETA-EYFP. During experiments, an optic fiber was positioned to target A1 and neuronal activity was recorded using a multichannel silicon probe in A1. B and C) Left: Blue light (430 nm) activates SOMs in SOM-Cre mice (B) or PVs in PV-Cre mice (C). Right: Raster plot showing a single neuron spiking response to 2.5ms pulses of blue light (blue shading). D) Diagram of tone trains and light pulses. Top: 2.5ms light pulses may occur at four time-points during each tone train repeat; before or after the train, or during the first or last tone pip. Bottom: close up of light pulse timing during each light-pulse time-point. E and F) PSTH of a single unit mean tone-evoked response with and without optogenetic activation of SOM (E) or PV (F) interneurons. The 2.5ms light pulse onset is 5ms after tone onset. Further analysis is drawn from the tone-response time window between 13 and 52ms (Green dashed lines). G-J) Scatter plots display the index of change (G and I) or difference (H and J) between light-on and light-off conditions for last versus first tone-evoked spiking responses within the tone-response time window. Each circle indicates a single neuron and its color indicates the significance of the change or difference between light conditions for each tone. Green asterisks above or below the unity line indicate a significant increase or decrease in population firing rates.

We recorded responses of excitatory and inhibitory neurons in awake mice presented with sequences of repeated trains, each composed of one of two chosen tone frequencies, while projecting light on AC either before the tone train, during the first tone, during the last tone, or after the tone train (Figure 6D). The timing of light activation was designed to elicit the volley of inhibitory activity just preceding the cortical response to the tone in excitatory neurons: the light pulse lasted from 5 to 7.5 ms after tone onset. This produced an elevated spiking activity in putative inhibitory neurons around 7-12 ms after tone onset, just before the typical tone-evoked activity, and suppressed tone-evoked activity (Supplementary Figure 4B, C). In some putative excitatory neurons, activating PVs or SOMs resulted in nearly full suppression of tone-evoked activity (Figure 6E, F). The FR elicited by the light pulse was not significantly different between the first and last tones for SOM-Cre mice (0.5±0.3Hz, p = 0.066, t(304) = 1.84), or PV-Cre mice (0.0±0.4, t(369) = 0.10) (Supplementary Figure 4D, E).

To estimate changes in connectivity between inhibitory and excitatory neurons, we compared the magnitude of change in excitatory neuronal firing on light-on and light-off trials before and after adaptation, for tone-evoked activity. We reasoned that if the synapses between inhibitory and excitatory neurons were facilitated after adaptation, eliciting the same number of spikes in inhibitory neurons should drive stronger changes in excitatory neuronal activity. Indeed, we found that briefly activating SOMs or PVs elicited relatively greater change in excitatory neuron activity during tone-evoked responses (SOM, 0.07±0.03, p = 0.0496, t(226) = 1.97; PV, 0.15±0.03, p = 1.5e-6, t(275) = 4.9) (Figure 6G, I). These findings suggest that the relative suppression from either inhibitory neuron becomes stronger after adaptation.

However, there were several caveats to this experiment. When we computed the firing rate difference rather than the relative index of change, we found that SOM and PV activation inhibited more spiking during the first tone compared to the last tone (SOM, -1.3±0.2 Hz, p = 2.7e-7, t(270) = -5.2; PV, -0.6±0.2 Hz, p = 5.2e-4, t(320) = -3.5) (Figure 6H, J). This result can be explained by the floor effect of near complete suppression of excitatory neuronal response: Because adaptation lowers the firing rate of excitatory neurons, inhibition cannot decrease the firing rate below 0, thus underestimating the amplitude of changes elicited by suppression of inhibition. Additionally, there was no significant difference in the relative effects of PV or SOM activation on the spontaneous firing rate – likely due to the low spontaneous firing rate of the neurons (SOM, 0.01±0.03, p = 0.743, t(170) = 0.32; PV, -0.00±0.04, p = 0.93, t(255) = -0.06) (Supplementary Figure 4F, H). As measured by the difference in firing rate, SOM and PV activation inhibited more spontaneous spikes before compared to after the tone train, but this likely due to reduced spontaneous activity following adaptation, similar to the effects during tone-evoked activity described above (SOM, -0.9±0.1 Hz, p = 9e-18, t(269) = -9.2; PV, -0.5±0.1, p = 2e-8, t(280) = -5.78) (Supplementary Figure 3G, I), likely due to floor effects. Combined, the results suggest that inhibition due to single spiked produced by either PVs and SOMs becomes relatively, but not absolutely, stronger with adaptation.

## Discussion

Our study dissected the complex roles of cortical inhibitory interneurons in adaptation to repeated sounds, identifying the differences in the contributions of two distinct interneuron subtypes, SOMs and PVs (8, 30). The roles of PV and SOM inhibition diverged sharply depending on the temporal context of tone presentation. We first showed that temporal adaptation was divisive across the frequency tuning curve of cortical neurons. Prior to adaptation, suppression of either SOMs or PVs evoked disinhibition of some and inhibition of other cortical neurons (Figure 3). In the adapted state, however, the effects of SOM suppression shifted to predominantly disinhibitory, whereas the effects of PV suppression remained heterogeneous (Figure 3D, F). These results demonstrate that SOMs are a major and unique contributor to adaptation in response to repeated stimuli. Furthermore, SOMs and PVs exhibited differential effects along the frequency tuning curve especially after adaptation: SOM suppression most strongly affected responses to preferred frequencies, whereas PV suppression only modulated responses to non-preferred frequencies (Figure 3G–J). These findings highlight the differences in the role of SOMs and PVs in controlling sound-evoked responses in the auditory cortex (Figure 7). Coupled with our finding that the magnitude of excitatory neuronal adaptation is highly correlated with the strength of inhibition by SOMs (Figure 5C), our results suggest that SOMs provide dynamically changing inhibitory drive to excitatory neurons dependent on the temporal dynamics of stimuli, whereas PV inhibitory effects are largely uncoupled from adaptation. Combined, the contribution of the inhibitory neurons maintains the frequency tuning of excitatory neurons after adaptation, resulting in divisive adaptation (Figure 1).

**Figure 7.**
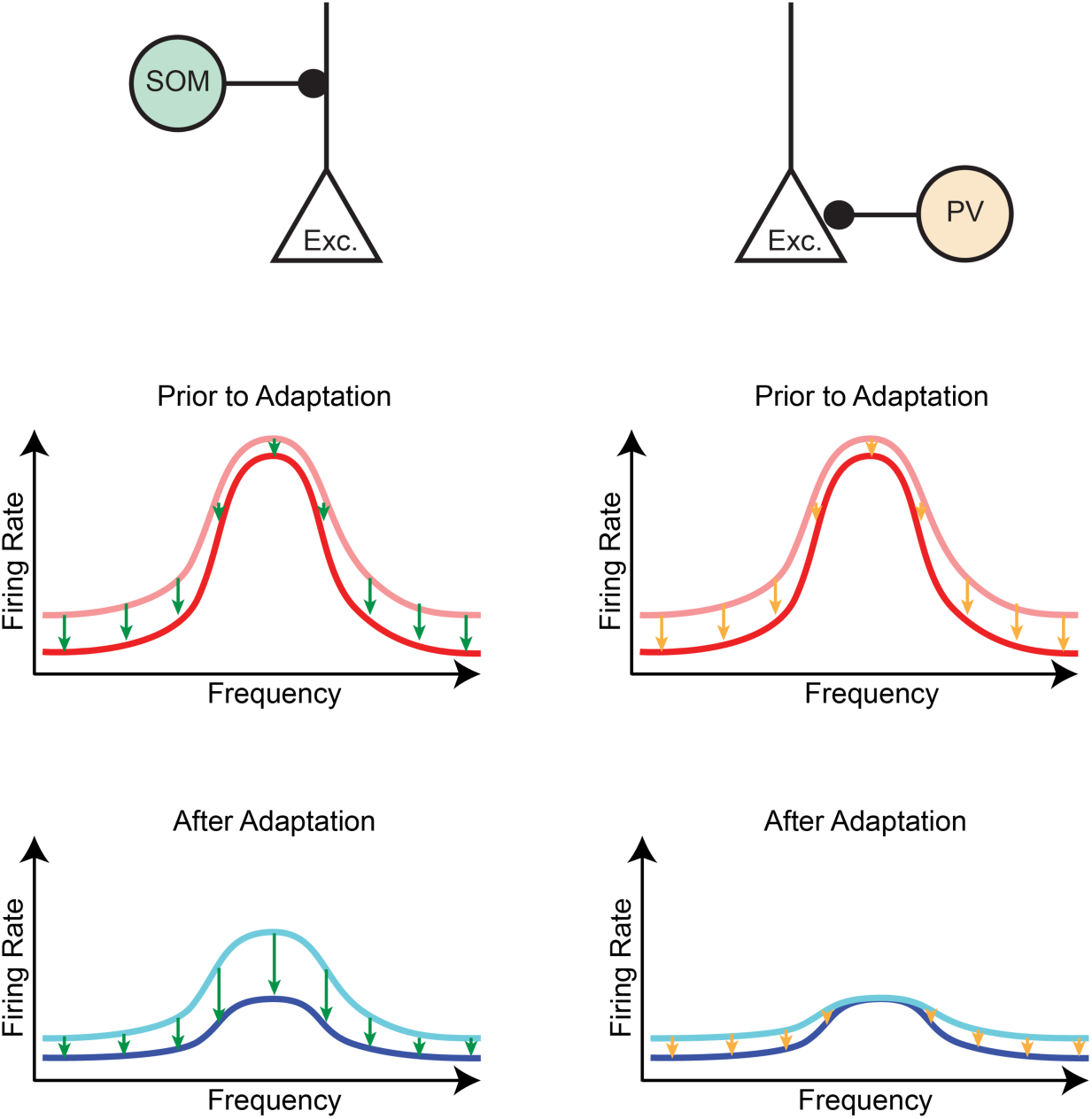
SOM inhibition reflects adaptation, while PV inhibition is insensitive to stable across adaptive regimes. Top: Circuit Diagram Middle: Model of typical pyramidal neuron frequency-response function in a non-adapted regime (red curve) as a combination of excitatory response (pink curve) and inhibitory suppression from SOM (green arrows) or PV interneurons (orange arrows). Bottom: Model of typical pyramidal neuron frequency-response function in an adapted regime (blue curve) as a combination of excitatory response (light blue curve) and inhibitory suppression (as above).

Cortical inhibitory interneurons are thought to play a crucial role in information processing and to shape how information is represented and transmitted within and between cortical neuronal populations (31). Molecularly distinct classes of interneurons are proposed to exert differential functions in information processing (1, 8, 32). Yet despite the differential morphology of the neurons, determining the differences in the action of PVs and SOMs in auditory cortical processing has proved elusive using static stimuli. Suppressing or activating PVs and SOMs during tone presentation led to a range of multiplicative and linear offset effects on excitatory tone-evoked responses (16–18). Similarly, measuring frequency tuning properties of PVs and SOMs did not reveal strong differences (33, 34). In the visual cortex, suppression or activation of PVs led to a linear or multiplicative shift on excitatory responses to oriented gratings (35–37), whereas modulation of SOMs led to differential responses to the center and surround (38). In the piriform cortex, suppressing SOMs led to uniform disinhibition of odor-evoked responses (39). This observed heterogeneity of effects is consistent with the bidirectional modulatory effects on tone-evoked responses that we observed (Figure 3, 4, Supplementary Figure 2). Such heterogeneity is hardly surprising given the complexity of cortical circuits. Multiple local inhibitory circuit pathways, such as direct monosynaptic inhibition or sign-reversing disynaptic disinhibition (3, 4, 40), may differentially influence pyramidal activity and contribute to these heterogeneous effects when an entire population of interneurons is selectively but incompletely suppressed. Furthermore, the interneuron classes of PVs and SOMs are themselves comprised of different subtypes, which furthermore exhibit different laminar distribution and connectivity patterns (*for recent reviews, see* 30, 41).

Interestingly, PV and SOM modulation produces consistent differences when the temporal dimension of responses is explored. SOMs respond to tones with a greater delay than PVs (33, 34). Thus, the effect of their modulation of excitatory neuronal activity is expected to follow a differential time course. Furthermore, SOMs, but not PVs, exert enhanced inhibition in accordance with behavioral habituation to stimuli over a time scale of days (28). In our study, we find that SOM-mediated inhibition, but not PV-mediated inhibition, is specific to temporal adaptation over a time scale of seconds. First, the effect of PV modulation is highly correlated from first to last tone pip, for both adaptive and non-adaptive neuronal responses (Supplementary Figure 3B, D). This means that PVs tend to exert the same type of modulatory effect across the timecourse of the tone train, suggesting that PVs are not affected by stimulus-history conditions. By contrast, SOM-mediated modulation effects on adaptive responses are more weakly correlated from first to last tone pip, and not correlated in non-adaptive responses (Supplementary Figure 3A, C). SOMs generally become more inhibitory after the tone train, suggesting that SOMs are strongly affected by stimulus-history conditions (Figure 3, 4). Additionally, the strength of SOM inhibitory influence on excitatory responses correlates with the magnitude of adaptation (Figure 5), i.e. SOMs more strongly inhibit neurons that adapt strongly. By contrast, PVs inhibit non-adapting neurons more strongly, at least during the first tone pip (Figure 3, 4). These results extend our previous findings in Natan et al. (19), which indicated that PVs merely amplify stimulus specific adaptation, while SOMs directly control it. Our present results demonstrate that SOMs dynamically control tone-evoked responses both across the spectral and temporal dimensions, whereas PVs provide a temporally uniform drive. These effects are likely due to their differential contribution to the inhibitory sidebands of excitatory neuronal receptive fields. The differential temporal and context-dependent effects may underlie the more fundamental distinction between the function of PVs and SOMs in sensory processing, which relies on long-term temporal integration.

What are the network mechanisms driving these differential changes in SOM inhibition during adaptation? A top candidate would be facilitation at the SOM to excitatory neuron synapse (42–45). We found that eliciting a brief burst of spikes in either PVs or SOMs predictably elicited suppression in excitatory neurons, which was relatively stronger after adaptation (Figure 6, Supplementary Figure 4). Combined, the results of this experiment and Natan et al., 2015, are consistent with the interpretation that synaptic strengthening between SOMs and excitatory neurons is likely to occur following adaptation.

However, multiple alternative mechanisms may underlie the observed phenomenon. First, repeated activation of facilitating synapses between SOMs and local pyramidal neurons may lead to increased inhibition relative to excitation that is specific to the stimulus, despite reduced spiking. Second, a distinct subpopulation of SOMs for which stimulus-evoked responses increase with repetition may entirely generate the observed increase in inhibition. Third, stimulus-specific reduced firing rates among SOMs may disinhibit PVs relative to pyramidal neurons, leading to increased inhibition among pyramidal neurons. Fourth, SOMs may relay a top-down or modulatory signal generated elsewhere that drives greater inhibition in specific stimulus contexts (46, 47). Testing each of these possibilities is required to disambiguate the specifics of the synaptic changes over the time scale of adaptation. Still, PV inhibition is unlikely to underlie the dynamics of any of these potential mechanisms due to their depressing synapses, predominant targeting of excitatory neurons, homogenous responses during adaptive stimuli and relatively low prevalence of direct top-down input (40). Thus SOM inhibition remains the primary candidate interneuron for generating the attenuation in cortical adaptation.

Some aspects of this study limit the extent to which the results may generalize. One limitation is that we only explored adaptation for a single temporal regime, with tones repeated at the rate of 2.5 Hz. Increasing the rate of tone presentation would potentially lead to a faster and stronger adaptation (48), which may recruit different intra-cortical circuits. Another limitation is that optogenetic manipulation did not allow for complete shutdown of interneuron activity (Figure 2H, I). Therefore, we measured the effect of modulatory changes and reduced dynamic range of evoked spiking from interneurons, rather than measuring the consequence of entirely removing a circuit element from the network. Future studies could extend these experiments focusing on the questions of adaptation dynamics by due to changes in inter-tone interval, timing and strength of optogenetic modulation.

Though it has been widely hypothesized that SOM and PV interneurons play different roles in cortical function, revealing those roles has been difficult, possibly due to a lack of consideration for temporal dynamics. In this study, we approached this problem by using stimulus parameters designed to induce adaptation over time and tuning. The results successfully define functional characteristics that differentiate SOMs versus PVs: SOMs appear to play a generalized role and likely underlie cortical adaptation to repeated stimuli, especially for well-tuned stimuli, while PVs do not. These findings support and add to our previous work showing that SOM inhibition underlies stimulus-specific adaptation (Natan et al., 2015), which itself has been confirmed by other groups (Hamm et al., 2016). Since adaptation is observed in every primary sensory cortex and is likely a canonical computation of all cortical regions, our results point the unique and generalized adaptive role SOM inhibition may play in canonical cortical circuitry.

## Materials and methods

### Animals

All experiments were performed in adult male mice (supplier - Jackson Laboratories; age, 12-15 weeks; weight, 22-32 g; PV-Cre mice, strain: B6;129P2-Pvalbtm1(cre)Arbr/J; SOM-Cre: Ssttm2.1(cre)Zjh/J) housed at 28° C on a 12 h light:dark cycle with water and food provided *ad libitum*. In PV-Cre mice, Cre recombinase (Cre) is expressed in parvalbumin-positive interneurons; in SOM-Cre mice, Cre is expressed in somatostatin-positive interneurons (49). This study was performed in strict accordance with the recommendations in the Guide for the Care and Use of Laboratory Animals of the National Institutes of Health. All of the animals were handled according to a protocol approved by the Institutional Animal Care and Use Committee of the University of Pennsylvania (Protocol Number: 803266). All surgery was performed under isoflurane anesthesia, and every effort was made to minimize suffering.

### Viral vectors

Modified AAVs encoding ArchT (AAV-CAG-FLEX-ArchT-GFP, UNC Vector Core) or ChETA (AAV-EF1a-DIO-ChETA-EYFP) were used for selective suppression or excitation, respectively, in PV-Cre or SOM-Cre mice.

### Virus injection

2-4 weeks prior to the start of experimental recordings, a 0.5 mm diameter craniotomy was drilled over primary auditory cortex (2.6 mm caudal and 4.1 mm lateral from bregma) under aseptic conditions while the mouse was anesthetized with isoflurane. A 750 nl bolus of AAV in water was injected into A1 (1 mm ventral from pia mater) using a stereotaxic syringe pump (Pump 11 Elite Nanomite, Havard Apparatus). The skull overlying A1 was thinned by gentle drilling. The craniotomy was covered with bone wax and a small custom head-post was secured to the skull with dental acrylic.

### Electrophysiological recordings

All recordings were carried out inside a double-walled acoustic isolation booth (Industrial Acoustics). Electrodes were targeted to A1 on the basis of stereotaxic coordinates and relation to blood vessels. In electrophysiological recordings, the location was confirmed by examining the click and tone pip responses of the recorded units for characteristic responses of neurons in A1, as described previously by our group in the rat (50–53) and by ours and other groups in the mouse (18, 19, 54–56). Mice were placed in the recording chamber, anesthetized with isoflurane, and the headpost secured to a custom base, immobilizing the head. Dental acrylic and bone wax was gently drilled away exposing auditory cortex, and a sillicon multi-channel probe (A1x32-tri-5mm-91-121-A32, Neuronexus) was slowly lowered to between 900 µl and 1100 mm into cortex, perpendicularly to the cortical surface. Electrophysiological data from 32 channels were bandpass filtered at 10-300 Hz for LFP and current-source density (CSD) analysis or at 600-6000 Hz for spike analysis, digitized at 32 kHz and stored for offline analysis (Neuralynx). Spikes belonging to single units were clustered using commercial software (Offline Sorter, Plexon) (50). Putative excitatory neurons were identified based on their expected response patterns to sounds and lack of significant suppression of the spontaneous firing rate due to light (33, 57).

### Acoustic stimulus

Stimuli were delivered via a magnetic speaker (Tucker-David Technologies), directed toward the mouse’s head. Speakers were calibrated prior to the experiments to ±3 dB over frequencies between 1 and 40 kHz, by placing a microphone (Brüel and Kjaer) in the location of the ear contralateral to the recorded A1 hemisphere, recording speaker output and filtering stimuli to compensate for acoustic aberrations (50). First, to measure tuning, a train of 50 pure tones of frequencies spaced logarithmically between 1 and 80 kHz, at 65 dB SPL relative to 20 µPa, in pseudo-random order, was presented 20 times. Each tone was 100 ms long (5 ms cosine squared ramped up and down) with an inter-stimulus interval (ISI) of 300 ms. Frequency response functions were calculated online for several multiunits. To construct the set of tone pip trains, 10 frequencies, spaced 0.13 octaves apart, were selected to cover a portion of multiunit frequency tuning curve from preferred to non-preferred frequencies. The tone frequencies were selected to overlap with the most neuronal frequency response areas, but did not exactly fall within the same frequency region of each neuron’s tuning curve. Each tone train consisted of 8 consecutive 100ms tone pips of the same frequency separated by 300 ms ISI. The frequency of each sequential train was pseudorandom and counterbalanced. Trains were separated by 2.4 seconds of silence.

### Light presentation

An optic fiber was use to direct 532 nm or 473 nm laser light (Shanghai Laser & Optics Century). After positioning the silicon probe, an optic fiber was placed over the surface of auditory cortex. To limit Becquerel effect artifacts due to light striking electrodes, we positioned the optical fiber parallel to the silicon probe (58, 59). For each train, light may be cast over AC to suppress interneurons during the first tone, the last tone, or during the silent period 400ms before or 400ms after the tone pip train. For ArchT mediated suppression, 532 nm light onset was 100 ms prior to tone onset, and lasted for 250 ms. For ChETA mediated excitation, 473 nm light onset was 5 ms after tone onset and lasted for 2.5 ms. At 180 mW/mm^2^, light pulses were intense enough to significantly modulate multiunit activity throughout all cortical layers.

### Immunohistochemistry

Brains were post-fixed in paraformaldehyde (4%, PFA) and cryoprotected in 30% sucrose. Coronal sections (50µm) for PV were cut using a cryostat (Leica CM1860) and SOM were cut using a vibratome (Vibratome 1000). Sections were washed in PBS containing 0.1% Triton X-100 (PBST; 3 washes, 5 min), incubated at room temperature in blocking solution (for PV 10% normal goat serum and 5% bovine serum albumin in PBST for 3hr; for SOM 1% normal horse serum with 0.1% bovine serum albumin and 0.5% Triton X-100 in PBS for 1hr), and then incubated in primary antibody diluted in blocking solution overnight at 4°C. The following primary antibodies were used: anti-PV (PV 25 rabbit polyclonal, 1:500, Swant) or anti-SOM (MAB354 rat monoclonal, 1:200, Millipore, clone YC7). After incubation, sections were washed in blocking solution (3 washes, 5 min), incubated for 2hr at room temperature with secondary antibodies (Alexa 594 goat anti-rabbit IgG 1:1000 for PV and Alexa 568 goat anti-rat IgG 1:750 for SOM), and then washed in PBS (3 washes, 5min each). Sections were mounted using fluoromount-G (Southern Biotech) and confocal images were acquired (Leica SP5).

### In vivo neuronal response analysis

#### Neuron-frequency pairs

Termed neuron-frequency pairs, each of the 10 frequencies used in the tone pip trains was tested separately for various trial-by-trail FR features for inclusion and exclusion from specific population analyses; tone evoked FR greater than 1 Hz, significantly greater tone-evoked responses compared to spontaneous activity, significant changes in FR in response to optogenetic illumination, significant differences between first and last tone-evoked FR.

#### FR analyses

Population PSTHs were calculated by first finding each neuron’s mean FR over pooled spike counts across all included frequencies, and normalizing by the mean peak FR in response to the first tone pip light-off condition, and then finding the mean across the neurons included in that population. Errorbars represent standard error. Tone-evoked FRs displayed in box and whisker plots were measured as the mean FR of each neuron between 10ms and 40 ms after tone onset. In scatter plots, each dot represents a neuron-frequency pair mean FR between 10 ms to 40ms after tone onset. Spontaneous activity before tone pip trains was measured from 400 ms to 350 ms before the first tone onset, and spontaneous activity after tone pip trains was measured from 400 ms to 450 ms after the last tone onset.

#### Linear fits across frequencies

Linear fits were calculated using linear regression (fitlm.m, Matlab) over 10 data points, one for each of the 10 frequencies tested. For each comparison condition (first or last tone pip response, with or without optogenetic suppression) the 10 data points were separately calculated as the mean FR over all repeats fitting those conditions. Linear fit error lines indicate standard error. Population average line was calculated as the mean of each lines y value across x from 0 to 1, and errorbars show standard error.

#### Statistical tests

Sign rank tests were used to test if first or last tone pip-evoked FRs or spontaneous FRs per trial were different within single neurons, and used for classification and population inclusion criteria. Rank sum tests were used to test FR differences between light conditions per trial within single neurons, and used for classification and population inclusion criteria. All population comparisons were tested using the student’s t-test. Correlations were measured across neuron-frequency pairs treated as separate samples in order to allow the dataset to reflect preferred and non-preferred frequencies. Since neuron-frequency pairs from the same neuron likely exhibit correlation, we used cluster-robust regression to compensate for a possible lack of independent sampling. Clustering samples by neuron adjusts the degrees of freedom, t-value, p-value, and corrected correlation coefficient *r*. In all figures and tests, single, double and triple stars indicate p < 0.05, 0.01 and 0.001 respectively.

## Acknowledgements

This work was supported by National Institutes of Health (Grant numbers NIH R03DC013660, NIH R01DC014700, NIH R01DC015527), Klingenstein Award in Neuroscience, Human Frontier in Science Foundation Young Investigator Award and the Pennsylvania Lions Club Hearing Research Fellowship to MGN. MNG is the recipient of the Burroughs Wellcome Award at the Scientific Interface. RGN was supported by National Institutes of Health (Grant number NIH NIMH T32MH017168).

